# Attraction, Dynamics, and Phase Transitions in Fire Ant Tower-Building

**DOI:** 10.1101/864306

**Authors:** Gary K. Nave, Nelson T. Mitchell, Jordan A. Chan Dick, Tyler Schuessler, Joaquin A. Lagarrigue, Orit Peleg

## Abstract

Many insect species, and even some vertebrates, assemble their bodies to form multi-functional materials that combine sensing, computation, and actuation. The tower-building behavior of red imported fire ants, *Solenopsis invicta*, presents a key example of this phenomenon of collective construction. While biological studies of collective construction focus on behavioral assays to measure the dynamics of formation and studies of swarm robotics focus on developing hardware that can assemble and interact, algorithms for designing such collective aggregations have been mostly overlooked. We address this gap by formulating an agent-based model for collective tower-building with a set of behavioral rules that incorporate local sensing of neighboring agents. We find that an attractive force makes tower building possible. Next, we explore the trade-offs between attraction and random motion to characterize the dynamics and phase transition of the tower building process. Lastly, we provide an optimization tool that may be used to design towers of specific shapes, mechanical loads, and dynamical properties such as mechanical stability and mobility of the center of mass.

## 1 Introduction

Collective aggregation is a prevalent behavior among social animals, where many individuals cluster together while feeding, defending against predators, or as a thermoregulation strategy, effectively reducing the exposed surface area per individual. Examples of species that aggregate include vertebrates such as penguins [Waters et al., 2012] and bats [Kerth, 2008, Roverud and Chappell, 1991] as well as insects such as beetles [Deneubourg et al., 1990], ants [Reynaert et al., 2006a, Theraulaz et al., 2002] and cockroaches [Ame et al., 2004, Jeanson et al., 2005]. While these aggregations are often planar, eusocial insects such as honey bees [Kastberger et al., 2011, Seeley, 2010], army ants [Franks, 1989], and fire ants [Mlot et al., 2011] extend this strategy and create three-dimensional assemblages. These self-assemblages are composed of a multitude of individuals who link their bodies, doing so without a global overseer and with limited cognitive abilities [Anderson et al., 2002].

The structures that these insects create are, in essence, autonomous materials that embed sensing, computation, and actuation. These properties are some of the long-standing aspirations in the fields of multi-functional materials and robotic materials [Hughes et al., 2019, Şahin, 2004]. Self-assembling agents have already begun to inspire robotic applications [Hamann, 2018]. For example, Del Dottore et al. [2018] have described the concept of “growing robots,” which are systems of a large number of individual robots working together to mimic biological growth in plants or groups of molecules or cells. Other collective robots are directly inspired by eusocial insects, such as the S-bots [Şahin et al., 2002, Groẞ et al., 2006], which form chains to collectively move larger payloads, just like ants working together to move larger food [Buffin and Pratt, 2016]. Also inspired by ants, Swissler and Rubenstein [2018] have developed robots with a new docking mechanism to form self-assembling structures. Another class of robots, inspired by termites [Werfel et al., 2014], build three-dimensional structures out of external building materials. Finally, the cube-shaped M-Blocks [Romanishin et al., 2015] construct aggregations out of their own bodies, using magnetism and angular momentum to climb on top of neighbors. These works represent examples from the emerging field of multi-agent robotic systems built out of many inexpensive individual robots and utilizing control strategies that may include redundancies to overcome individual malfunctions. While much of the focus in robotics has been on developing the hardware, the algorithmic development of assembling processes has often been overlooked. We address this gap by borrowing tools from computational material science and characterize the dynamics of 3-dimensional aggregation formation inspired by fire ant towers.

In nature, red imported fire ant (*Solenopsis invicta*) towers tend to occur in the event of flooding. Initially, fire ants gather together to form hydrophobic rafts [Mlot et al., 2011, 2012] to float above the water surface. When the rafts approach vegetation emerging from the surface, they may attach to the vegetation and form towers on top of their floating rafts. In a recent study, Phonekeo et al. [2017] described an experimental assay of the tower-building process in fire ants. In their analysis, the authors propose four rules which allow ants to build towers:

1. Do not move if ants are on top of you.
ii. If atop other ants, repeatedly move a short distance in a random direction.
iii. Upon reaching available space adjacent to non-moving ants, stop and link with them.
iv. The top layer of the tower is not stable unless there is a complete innermost ring of ants gripping each other around the rod.

The work of Phonekeo et al. [2017] shows an agreement between the resulting tower shapes in the long-timescale limit; however, it does not explore the time dynamics and parameter space systematically. This is what the present work aims to do, since local rules such as these provide a systematic way of analyzing collective behavior through agent-based modeling, and importantly, they are directly implementable in swarm robotic systems. By simulating the behavior of individuals following a set of local rules, it is possible to investigate how local interactions between agents lead to global emergent behavior and explore the space of possible behavior beyond what is possible with experiments.

Modeling efforts of collective behavior using local behavioral rules include the boids model (or Vicsek model) [Reynolds, 1987, Vicsek et al., 1995], which simulates agents moving under attraction, repulsion, and alignment. However, it best describes the behavior of more sparse collectives such as flocks of birds or schools of fish. To model ants building a tower, we must account for dense aggregations where the interaction range is limited to a short length scale, preferably defined by the size of an individual agent. Models of more dense collective assemblies include aggregation in slime mold based on chemical signal amplification [Levine et al., 1997, Umeda and Inouye, 1999], and nest building in wasps using an agent-based model in which swarms of builders deposit bricks and build up a nest [Bonabeau et al., 2000, Theraulaz and Bonabeau, 1995]. However, we must consider moving ants, which climb the tower and form the shape, as well as stationary ants that support the structure. Based on the similarity to the aggregation of inanimate systems such colloids [Deneubourg et al., 2002, Vernerey et al., 2018], we reason that ant tower building would experience dynamic phase separation process including nucleation [Vlasov, 2019], jamming [Bak, 1996], and ripening [Voorhees, 1985]. These phase transitions are also observed at the thermodynamic transition between phases of matter, which have been studied experimentally [Panagiotou et al., 1984] as well as computationally [Navarro and Fielding, 2015, Rovere et al., 1990]. Hence, we formulate an agent-based model with a set of behavioral rules that lead to aggregation and experience dynamical phase transitions.

Section 2 describes the details of the model we study in the present work and lays out the modifications to the local rules (presented above) that we introduce to achieve tower-building. In Section 3, we explore the parameter space of the local rules to identify the impacts of each component: locking, unlocking, and attraction. We find that towers undergo a phase transition when varying the attraction parameter, and explore how this phase transition changes across various densities. Finally, we introduce an optimization algorithm to generate the largest possible tower for a given density of agents in the system. In Section 4, we discuss the significance of the results and talk about implications for both the understanding of collective biological systems and the design of multi-agent robotic control strategies.

**Figure 1:**
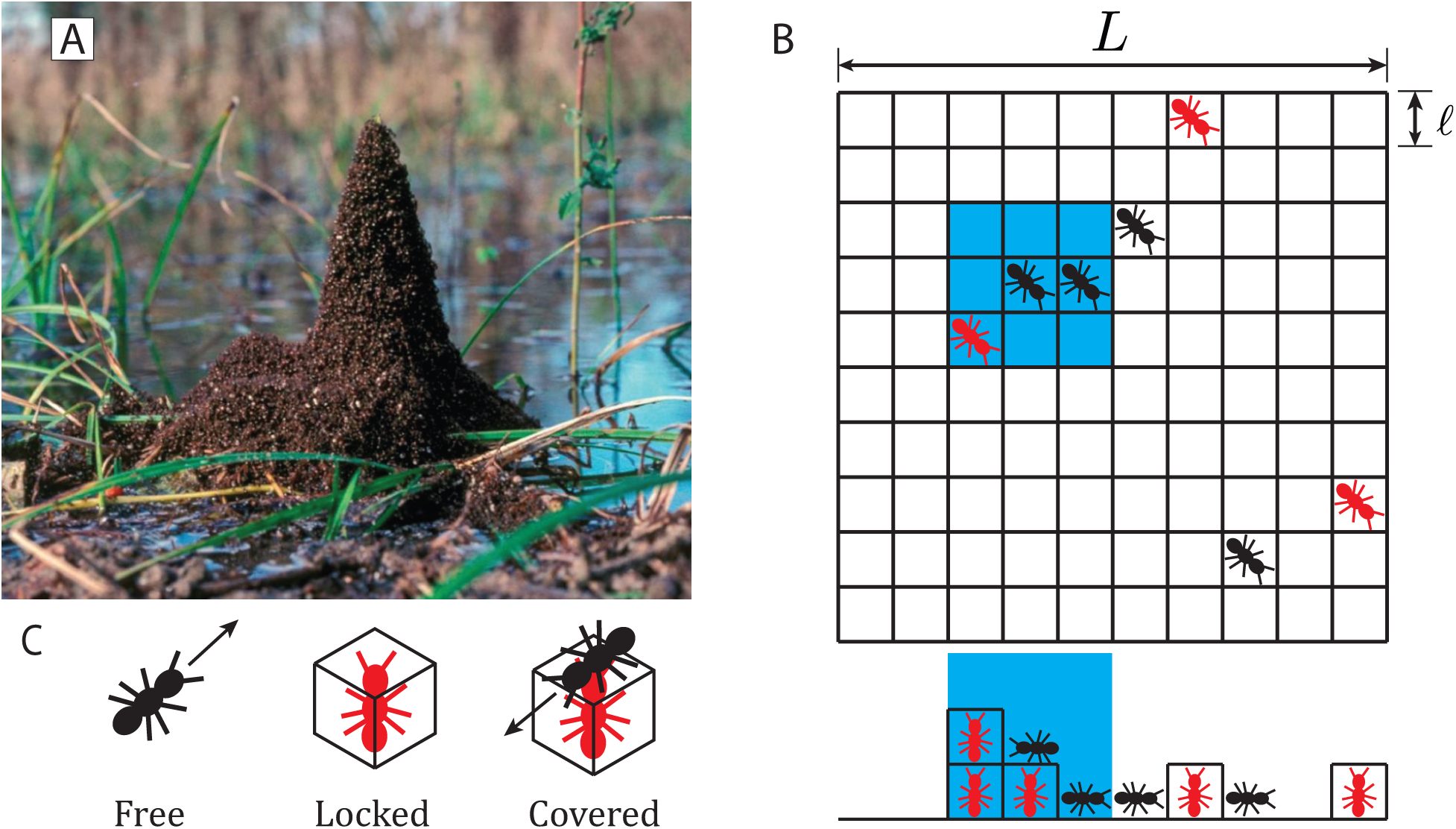
(A) Formation of towers by fire ants around vegetation, reproduced from Phonekeo et al. [2017]. (B) Schematic of computational model considered in this study. Individual free agents (black) move to an adjacent square at every time step, while locked agents (red) remain stationary. The side view below the grid shows the 3-dimensionality of the model arena. The blue region represents the neighborhood of adjacent locations which an agent considers when searching for locked neighbors and for determining its attractive force. (C) Schematic representation of the three states an agent can assume in the present model. Free agents move with constant velocity, locked agents stop to build towers, while covered agents cannot unlock.

## 2 Agent-based model

We consider a system of *N* individual agents simulated to move in a *L × L* × ∞ arena, discretized into a cubic lattice made of voxels of volume *ℓ × ℓ × ℓ*. The volume of an individual agent is set to the be volume of a voxel, where *ℓ* ≡ 1. At each time step, an individual agent can move into one of its 26 neighboring voxels: 9 above, 9 below, and 8 on the same level. A schematic of agents moving within the arena is shown in Figure 1B. In the present work, we will not consider the effects of solid wall boundaries and will instead implement periodic boundaries. The horizontal plane of the arena, therefore, contains periodic boundary conditions - when an agent leaves the right side of the arena, for example, it re-enters the left side. Periodic boundaries are also taken into account when distances between agents are calculated. The equations that define the periodic boundary conditions are given in (5) and (6) in Appendix B. The vertical direction of the arena is semi-infinite, extending upward from a solid floor.

Agents move horizontally and climb if they the voxel they intended to move into is occupied by a locked agent. Note that the local rules described above, from Phonekeo et al. [2017], refer to agents “linking” with one another, while in the present work we will refer to an agent that stops to support tower building as “locked.” Each pixel along the horizontal plane has an associated height equal to the number of locked agents on top of each other in that location. The free agents, therefore, are moving under 2-dimensional rules along the surface defined by locked agents, which is embedded in 3-dimensions. If an agent attempts to climb on top of neighbors to a voxel that is more than *ℓ* higher, it does not move at this time step.

Agents in the model may take on three different states, depicted in Figure 1C: free, locked, or covered. A free agent may move around the arena according to a specific set of behavioral rules with a constant velocity of one voxel per time step. Movement order is chosen randomly at each time step. To prevent two individuals from occupying the same position, if two free agents move to the same voxel, the second agent to arrive randomly chooses an unoccupied voxel adjacent to the target voxel. Locked agents are those which have decided to become a part of a tower and allow their neighbors to climb on top of them. We explore different schemes for the decision to ‘lock’ as defined in Sections 2.1 and 2.2. Covered agents are locked agents with at least one other agent on top of them. Each time step consists of first evaluating movement for all individuals and then evaluating locking decisions for all individuals based on their new configuration. We will not consider the effect of stability and assume that each agent has infinite strength to support neighbors.

It is likely that pheromones play a role in fire ant tower building, but for the present study, we consider whether this behavior can arise from solely physical proximity to neighbors. Hence, an agent can sense which of its surroundings 26 voxels are occupied by another agent. This local model will allow for easier implementation by collective robotic systems, as it merely requires local sensing.

Unless specified otherwise, all simulations contain *N* = 1, 000 agents moving in a 100 × 100 × ∞ arena, corresponding to a density of 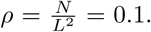 Based on preliminary simulations, we have chosen to evaluate each trial for 500,000 time steps, for which 97.8% of all simulations considered reached a steady state, where the largest tower size remained approximately constant (± 5% of *N*) for at least the last 100,000 time steps of the simulation. Exceptions will be discussed below in Section 3.2.

### 2.1 Diffusion-limited aggregation

We start by investigating whether a dynamic simulation of the proposed local rules above can lead to tower-building. As we are not considering effects of stability, we will ignore rule (iv) in the present study. We simulate the rules (i)-(iii) from Mlot et al. [2012] and Phonekeo et al. [2017] with a naive approach to what constitutes an available space adjacent to non-moving agents, assuming no direct knowledge of the agents about where they are relative to the rest of the tower. At each time step, each individual agent randomly chooses an adjacent square to move into, performing a random walk and fulfilling rule (ii). When an agent arrives in a voxel with at least one locked neighbor sharing a corner, edge, or side, it decides to lock, fulfilling rule (iii). Locked agents remain in place, and allow others to move on top of them. Finally, when agents climb on top of locked agents, the locked agent’s status changes to covered, fulfilling rule (i).

This model leads to aggregations which grow horizontally rather than upward. An example of a final configurations from one such simulation is shown in Appendix A and correspond to the boxed-in panel of Figure A1. This is illustrated in supplemental video S1, where each tower growing outward in fractal shapes from a center point. This behavior arises due the higher likelihood of an agent performing a random walk to find other agents near the outer edge of the aggregation.

These results closely resemble a phenomenon known as diffusion-limited aggregation (DLA) [Witten and Sander, 1981]. DLA was developed to model the aggregation of metal particles which gather in wispy, fractal shapes, similar to the simulated agent aggregation in Figure A1 for *P*_*u*_ = 0, *k*_*nl*_ = 1. DLA has also been observed in experimental colloidal aggregation systems, as in Reynaert et al. [2006b]. Without any rule modifications, DLA is unable to form dense aggregations of agents, because agents on the edge of the aggregation shadow those closer to the center. Hence, we propose several modifications to the behavioral rules which are necessary to mimic the time dynamics of tower shapes experimentally observed by Phonekeo et al. [2017].

### 2.2 Rule modifications to achieve tower-building

#### 2.2.1 Probability of unlocking

First, we allow locked agents to unlock with a constant probability, as long as they are not covered by other. This allows individuals past the first locked neighbor they encounter and move further in toward the center of an aggregation. To model this, we introduce a constant probability of unlocking *P*_*u*_ which applies equally to all uncovered locked agents. This rule introduces a distinction between locked agents and covered agents - covered agents cannot unlock.

#### 2.2.2 Neighbor-influenced locking probability

Second, we loosen the requirement that agents must lock upon encountering another locked agent, and instead allow for their probability of locking to increase with an increasing number of locked neighbors. This new rule (ii) replaces the previously discussed rule that individuals lock immediately upon finding a locked neighbor. Instead, an individual has a probability to lock based on the number of locked agents in its neighborhood. We define this probability of neighbor-influenced locking as *P*_*nl*_ = *k*_*nl*_*N*_*n*_, with *N*_*n*_ representing the number of locked agents in an individual’s neighborhood and *k*_*nl*_ specifying the increase in probability for each additional neighbor. The neighborhood is defined as a distance of one above below, or horizontally adjacent to the agent’s location, highlighted by the blue region in Figure 1B.

The overall probability that a free agent chooses to lock is given by,

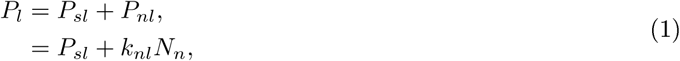

where *P*_*sl*_ is a probability of spontaneously locking. The probability of spontaneous locking provides a baseline probability of locking, to allow for individuals to randomly seed towers. In our simulations, we keep this probability small and set it to 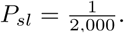 The neighbor-influenced locking factor provides the urgency with which an agent locks next to locked neighbors.

#### 2.2.3 Attraction forces

As we show below in Section 3.1 and Appendix A, the two rule modifications above are unable to reproduce large tower-like structures. Therefore, we extend the random walk model discussed above, and add an attractive “force” representing a behavioral tendency to cluster together. Under this effect, individual agents search their immediate local neighborhood for other agents, and move toward the center of all neighbors. This motion is then perturbed by the randomness associated with a simple random walk model. The resulting velocity is given by,

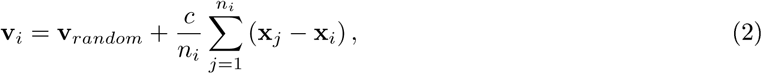

where *n*_*i*_ is the number of neighbors in the agent’s immediate neighborhood sharing at least one corner, edge, or side with the agent’s current position, and *c* is ratio of the magnitude of attraction relative to the magnitude of randomness. Each agent moves toward the available voxel most closely aligned with the direction of **v**_*i*_. The resulting normalized velocity, 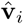 is defined in (7) in Appendix B. Each agent moves into the voxel defined by the surface height at the resulting pixel.

### 2.3 Set of modified behavioral rules

With the three modifications mentioned above, we modify the first three rules of Phonekeo et al. [2017] and Mlot et al. [2011] into four new local rules:

1. Do not move if agents are on top of you.
ii. Upon reaching available space adjacent to non-moving agents, stop and lock with them with probability *P*_*l*_ = *k*_*nl*_*N*_*n*_.
iii. If stopped and locked, but uncovered by other agents, spontaneously unlock with probability *P*_*u*_.
iv. If free to move, move generally toward your neighbors with some random noise as defined by equation (2).

### 2.4 Measurements of tower geometry

Each simulation is post-processed to measure the geometry of each tower in order to determine how tower-like the aggregation is. For the final configuration of each simulation, a 2-dimensional height map is constructed by assigning each pixel in the 2D projection of the arena a value equal to its maximum height (Figure 2A). We treat the resulting *L × L* array of pixel values as an image and apply connected-component analysis [Shapiro, 1996] to identify different towers - a labeled image is generated where any two non-zero pixels that share a corner or edge have the same label. Each agent in the simulation is then given the label corresponding to its horizontal position within the labeled image. As we are interested in building a single large tower, properties for the tower containing the largest number of agents from each simulation are reported. Three tower properties are considered: number of individuals per tower, maximum tower height, and the ratio of the tower height to its equivalent diameter. Equivalent diameter is defined as the diameter of a circle with area equivalent to the tower’s base (Figure 2A).

## 3 Results

To gain an intuition for the effects of the modifications to the tower-building rules discussed in Section 2.2, simulations were run over a range of locking and unlocking parameters, *k*_*nl*_ and *P*_*u*_, across multiple attraction parameters, *c*, and, in Section 3.3, across varying densities of agents in the system, *ρ*. We begin with a parameter sweep across the locking and unlocking parameters and attraction parameter in Section 3.1. Then, selecting a pair of locking and unlocking parameters, we systematically vary attraction *c* to show a rapid phase transition, and investigate the time dynamics of tower properties, both near and far from the phase transition in Section 3.2. In Section 3.3, we vary the density of agents along with attraction, and observe, in Section 3.4, that the center of mass of the towers may continue to move. Finally, we optimize for tower size and height in Section 3.5 and identified a set of parameters where a tower formed of nearly all individuals in the simulation.

### 3.1 Tower geometry

To explore the range of possible tower shapes in the model, we sweep the parameter space of the three rule modifications, including probability of unlocking *P*_*u*_, neighbor-locking factor *k*_*nl*_, and attraction factor *c*. Resulting tower properties and example final configurations from these simulations are shown in Figure 2. Every data point represents the mean of the largest tower’s properties for each of ten simulations. The left column of the array of tower properties, representing simulations with *c* = 0, shows that without attraction, towers tend to contain a small number of agents, a small height, and an especially low height-diameter ratio. These simulations with *c* = 0 represent the first two rule modifications - individual unlocking and neighbor-influenced locking - alone. From the measured tower properties in Figure 2B, we see the effects of the first two rule modifications without attraction. The aggregations with the largest number of agents are found in the simulations with parameters *k*_*nl*_ = 1 and *P*_*u*_ = 0, representing the case of no rule modifications at all. These aggregations lead to diffusion-limited aggregation as discussed above and shown in Supplemental video S1. The locking and unlocking rule modifications, therefore, decrease the number of agents in the largest aggregation. They do provide an increase in tower height and the height-diameter ratio. This increase is modest, however, with the tallest average tower height reaching 3.4 agents tall for *P*_*u*_ = 0.002, 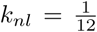, corresponding to a height-diameter ratio of 0.314. The largest height-diameter ratio occurs for the parameters *P*_*u*_ = 0.02, *k*_*nl*_ = 1, reaching a value of 0.49, with a corresponding average height of 2.2 and 19.9 agents in the largest tower for each simulation. Finally, the simulations with *c* = 0 and *P*_*u*_ = 0.2 with a small lock factor 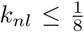 finish the simulations without forming aggregations. Supplemental video S2 and the *c* = 0 configuration snapshot in Figure 2C show the dynamics and final configuration, respectively, of one such simulation which is unable to form aggregations, with parameters *P*_*u*_ = 0.2, 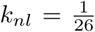, *c* = 0. These tower measurements show that without attraction, all of the tested parameter sets produce aggregations that remain small in number of individuals, do not reach average heights more than 3.4 layers, and remain wide and shallow.

**Figure 2:**
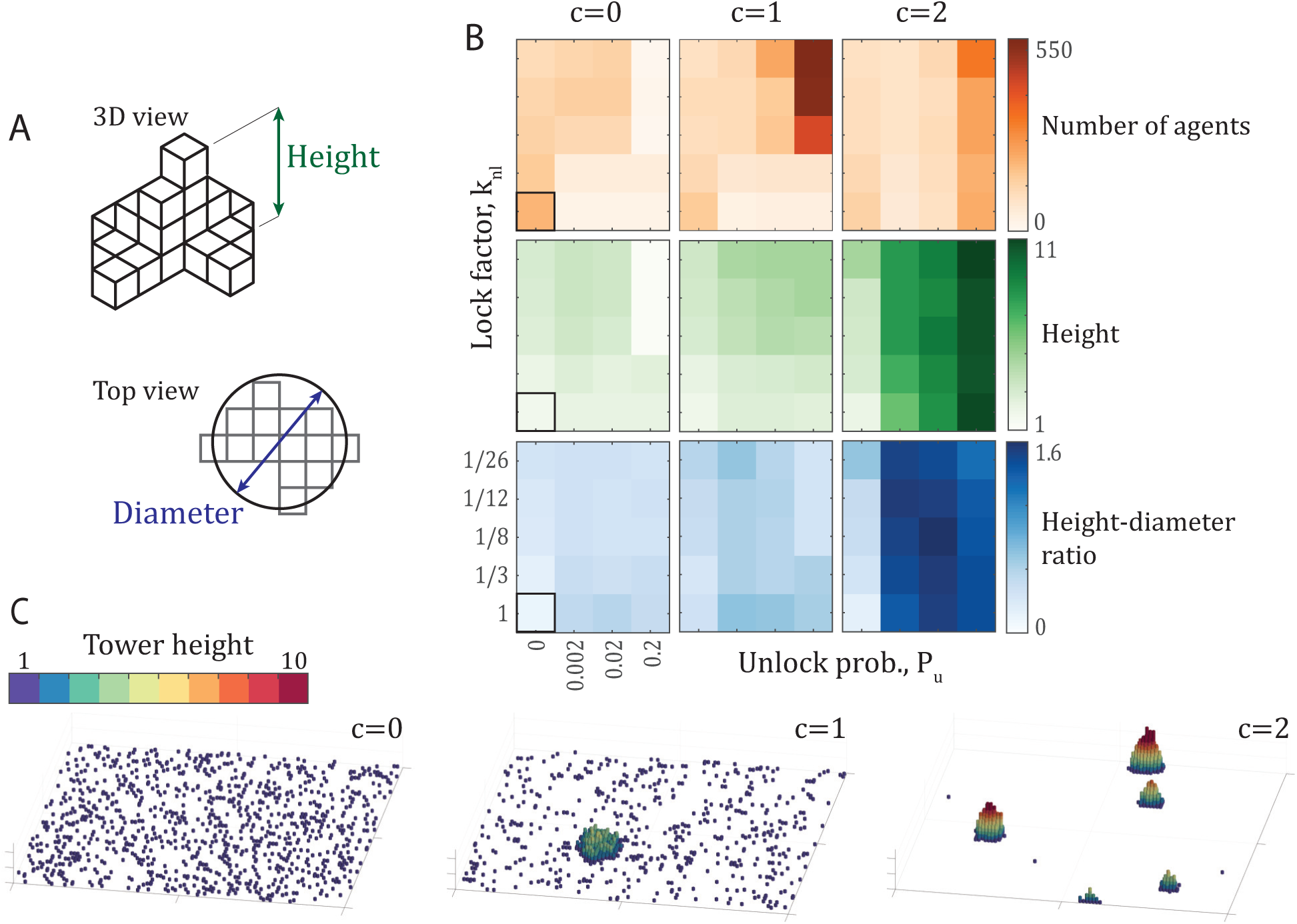
(A) Schematics of tower height and diameter, defined in Section 2.4. (B) Average number of agents (orange), tower height (green), and aspect ratio (blue) of the largest tower across a variety of parameters. Every data point represents the mean of the properties of the largest tower from each of ten simulations after 500,000 time steps. Aspect ratio (blue) is defined as the ratio of tower height to the diameter of a circle with area equal to the tower base. The bordered square represents the case of no rule modifications, with parameters *P*_*u*_ = 0, *k*_*nl*_ = 1, *c* = 0. Note that the axes of the property comparisons are not linear. (C) Examples of the final configuration of agents after 500,000 time steps for *c* = {0, 1, 2} with *P*_*u*_ = 0.2, 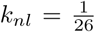. Each panel shows the entire 100 × 100 arena. The videos of the simulations that result in these final configurations are shared as supplemental videos S2-S4.

When an attractive force is added, larger aggregations form, as shown by the center column of Figure 2B for an attraction ratio of *c* = 1. As unlock probability *P*_*u*_ increases and lock factor *k*_*nl*_ decreases, larger aggregations form, with the largest reaching over 500 individuals. However, these largest aggregations have the smallest height-diameter ratios of this set, showing that these large aggregations are particularly wide, as is visible in the *c* = 1 example in Figure 2C and supplemental video S3. Increasing the attraction ratio to *c* = 2 finally reveals a more typical tower-like shape, with taller aggregations, even reaching a height of 11 agents. Interestingly, these taller towers contain fewer agents than the *c* = 1 case. The reason for this is clear in the snapshots shown in Figure 2C and supplemental video S4-stronger attraction yields more densely-packed towers with larger height-diameter ratios - the towers are so dense that multiple, smaller towers form instead of most individuals aggregating into a single tower.

### 3.2 Phase transition and time dynamics of tower-building

The example configurations shown in Figure 2C represent the same set of locking and unlocking parameters, *P*_*u*_ = 0.2, 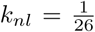 across *c* = {0, 1, 2}. These locking and unlocking parameters give the largest towers for both *c* = 1 and *c* = 2, but no aggregations at all for *c* = 0. To investigate the effects of the attraction ratio *c* further, we selected a fixed pair of locking and unlocking parameters and explored both the height and number of agents in the largest tower in the system for a densely sampled range of the attraction parameter *c*. The results of these simulations are shown in Figure 3A and 3C. The presence of a phase transition occurs between *c* = 0.92 to *c* = 1.06, where the number of agents in the largest tower climbs from close to 0 to over 700 agents. The results show that as *c* increases beyond that critical value, the number of individuals in the largest tower decreases (Figure 3A) while the height of the largest tower increases (Figure 3C).

In Figure 3B, we show the time dynamics of the number of agents for tower for two cases, close to the phase transition and further from it. To illustrate tower growth further from the phase transition, Figure 3B shows the time histories of all 10 simulations for *P*_*u*_ = 0.2, 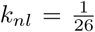, c = 2.0 in green and the mean of all simulations in black. One of these simulations is shown in supplemental video S4. The tower formation process in this model demonstrates two time scales: the time scale of initial nucleation, and the time scale of growth. Nucleation generally occurs within the first 5,000 time steps, the first 1% of each simulation. After nucleation, towers often continue to grow slowly through the rest of the simulation. Occasionally, two towers will merge into one, which manifests as a sharp jump in the time histories of Figure 3B. Some of these tower collisions last through the rest of the simulation, while others briefly merge and then separate again, which shows up as a sharp peak in the time history of tower size. The fast nucleation followed by slow growth seen for *c* = 2.0 is typical for most simulations in the present work.

However, there are some examples, particularly within the phase transition regime, for which a critical slowing down occurs. Trajectories close to the phase transition are shown in Figure 3D. Two trajectories are shown for each of *c* = {0:96; 1:0; 1:3} with *P*_*u*_ = 0.2, 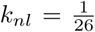. The critical slowing down is particularly evident for the *c* = 0.96 trajectories, where agents aggregate into a tower after 250,000 time steps while the other never transitions out of the disordered state. The *c* = 1.0 trajectories also show variation in nucleation time, although in this case, all simulations have transitioned to their aggregated state, in which the largest tower contains at least 100 agents. The variation in tower size is highest for these examples, varying in size by 200 or more individuals. There are also cases where the towers continue to grow in size, even after 500,000 time steps, which can also be seen in the case of *c* = 1.3.

**Figure 3:**
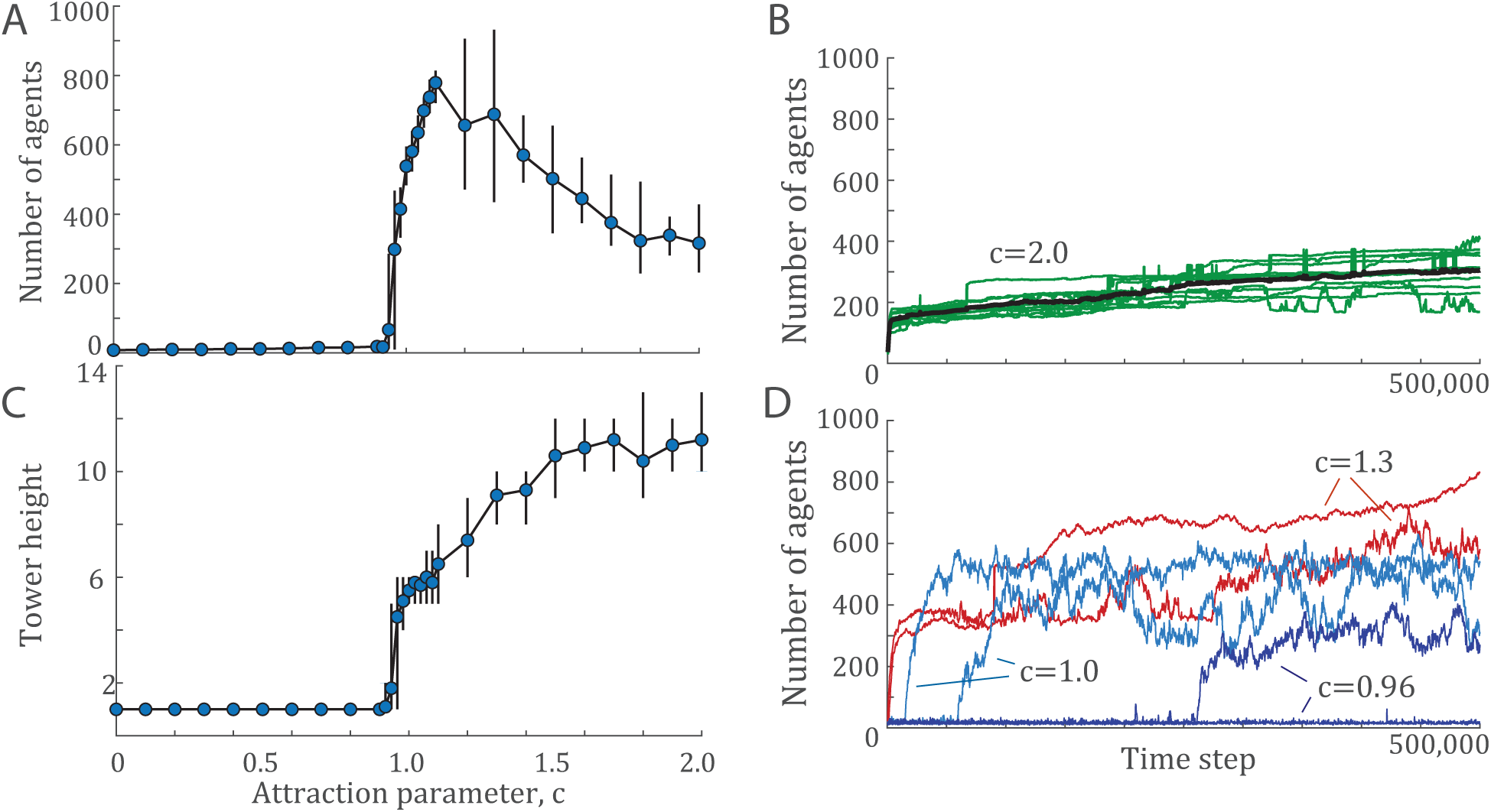
(A) The number of agents and (C) tower height of the largest tower across a range of attraction parameters, *c* for a given pair of locking and unlocking values, *P*_*u*_ = 0.2, 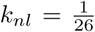. Each data point shows the mean value over ten simulations of 500,000 time steps each. Error bars indicate the maximum and minimum values observed. (B) Example time histories from ten simulations of *P*_*u*_ = 0.2, 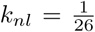, *c* = 2.0 in green with the mean at each time step shown in black. (D) Time histories from two examples for several *c* values near the phase transition, *c* = {0.96, 1.0, 1.3}.

As discussed in Section 2, the simulation time of 500,000 was chosen because nearly all simulations have reached a steady state. The cases highlighted in Figure 3D represent the exceptions, and there is no guarantee that these simulations will ever converge. The figure shows that *c* = {0.96, 0.98} are the only cases that give a mixture of aggregated and non-aggregated results.

### 3.3 The effect of density

The parameters varied up to this point in the model represent entirely behavioral parameters, that is, those associated with the decision-making of individuals. While these parameters are testable within multi-agent robotic examples, they do not represent a variable that can be systematically changed in experiments with live fire ants, or robots, in order to test the predictions of the model. To develop a set of testable predictions, we turn to explore the parameter of density of agents, *ρ*.

In our model, density is varied by changing the number of individuals in a fixed arena size. The computational complexity of the model is 𝓞(*N*^2^), so practical limits of computational time place an upper bound on density we explore here. In a 100 × 100 × ∞ arena, our test set is *N* = {200, 500, 750, 1000, 1500, 2000} which corresponds to the densities *ρ* = {0.02, 0.05, 0.75, 0.1, 0.15, 0.2}. We will use unlocking and locking parameters of *P*_*u*_ = 0.2, 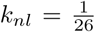 for consistency with Section 3.2.

The results of these simulations are presented in Figure 4, showing several key differences and similarities across densities. As density increases, less attraction is required for tower formation. The data points highlighted by circles in Figure 4B show the critical attraction ratio *c*^∗^, which represents the minimum value of *c* for which the largest aggregation is at least 100 individuals, representing the onset of the phase transition. This result also implies that there exists a critical density across a range of attraction factors, below which no tower formation occurs. Another key result is that the largest towers, in terms of number of agents per tower, occur shortly after the transition from no towers at all. Beyond this point, tower height and ratio increase while number of agents decreases. Finally, tower shape remains close to constant across densities, particularly in height-diameter ratio.

### 3.4 Moving towers

The introduction of unlocking probability effectively adds noise to the system (equivalent to higher temperatures in thermodynamic systems), which allows towers to move. Agents locking on one side of the tower while others unlock on the other side can lead to tower motion. Traces of the center of area for each tower from two example simulations may be seen in Figure 5B. To quantify this phenomenon, we consider the motion of towers as a Brownian random walk and investigate the diffusion coefficient of each tower. The diffusion coefficient (*D*) for a Brownian random walk follows the relationship,

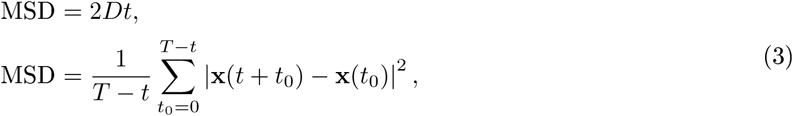

for each trajectory of length *T*. Therefore, we measure the mean square displacement (MSD) of each tower in each simulation over a variety of times, *t* = {0, 250, 500,…, 12, 500}, and perform a linear fit for each tower trajectory. The average slope of these lines is then twice the diffusion coefficient (Figure 5A.).

These results show that the maximum diffusion occurs in the highest density regime, and for the lowest attraction parameters that generate aggregations, particularly for *ρ* = 0.2 at *c* = 0.75 and *c* = 1.0. These towers have lower height-diameter ratios, as seen in Figure 4B, which leads to a larger proportion of agents on the surface of the tower, and therefore a higher probability that individuals on the surface will be unlocking. The towers at *c* = 0.75 have a smaller number of agents than those of *c* = 1.0, which leads to an even higher proportion of individuals on the surface. This is illustrated in Figure 5C, showing the time-evolution of tower configurations for two example simulations (see also supplementary videos S5 and S6).

**Figure 4:**
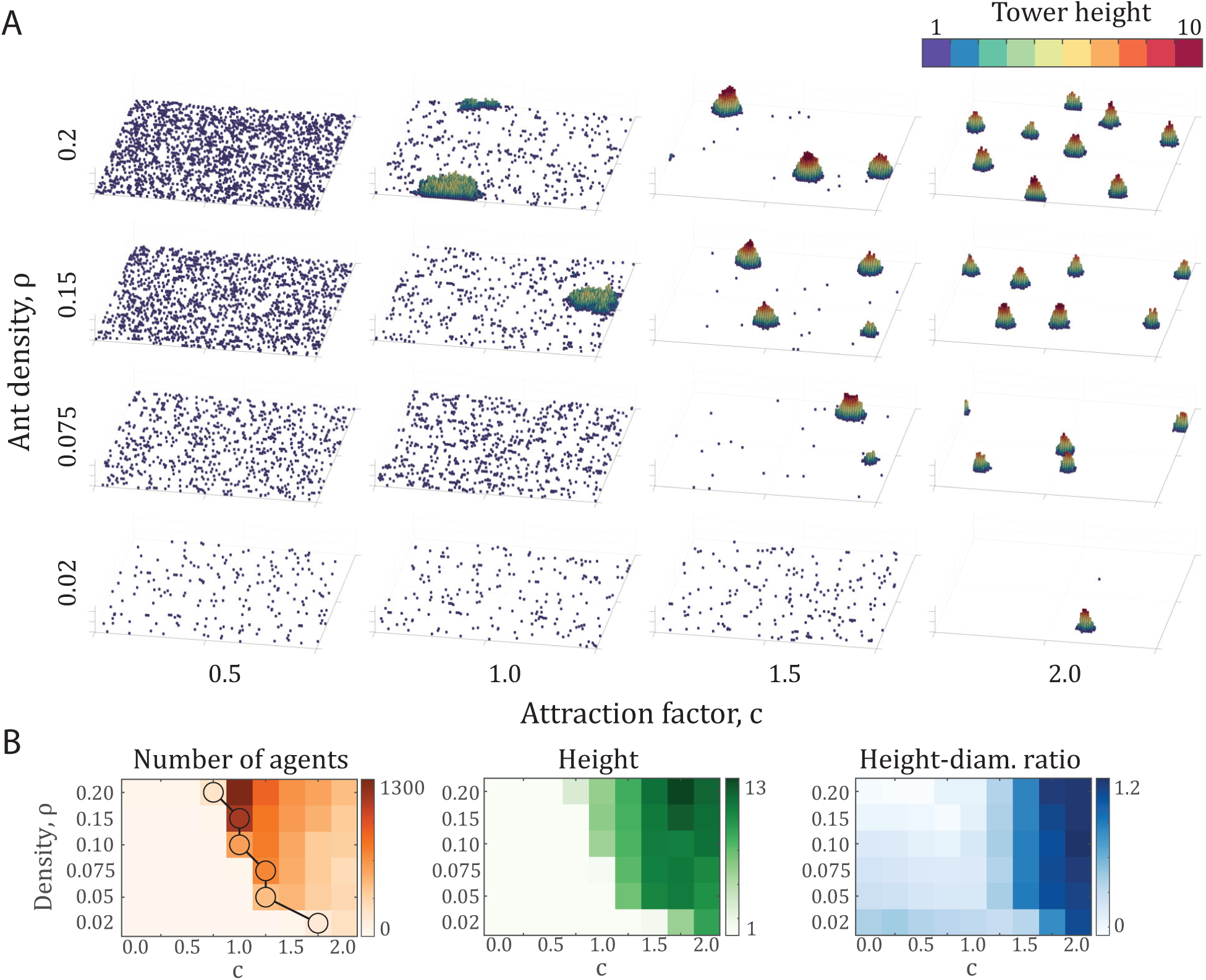
(A) Final configurations of various simulations after 500,000 time steps, comprising densities of *ρ* = {0.02, 0.075, 0.15, 0.2} and attraction factors of *c* = {0.5, 1.0, 1.5, 2.0}. Each panel shows the entire 100 × 100 arena. (B) Number of agents, height, and height-diameter ratio over a range of density *ρ* and attraction factor *c*. Note that the vertical axis is not linear. Each data point represents the average properties of the largest tower from ten simulations after 500,000 time steps. The circles on the density plot represent the onset of phase transition. The circles indicate the minimum attraction coefficient *c* for each density at which the largest tower contains at least 100 agents.

**Figure 5:**
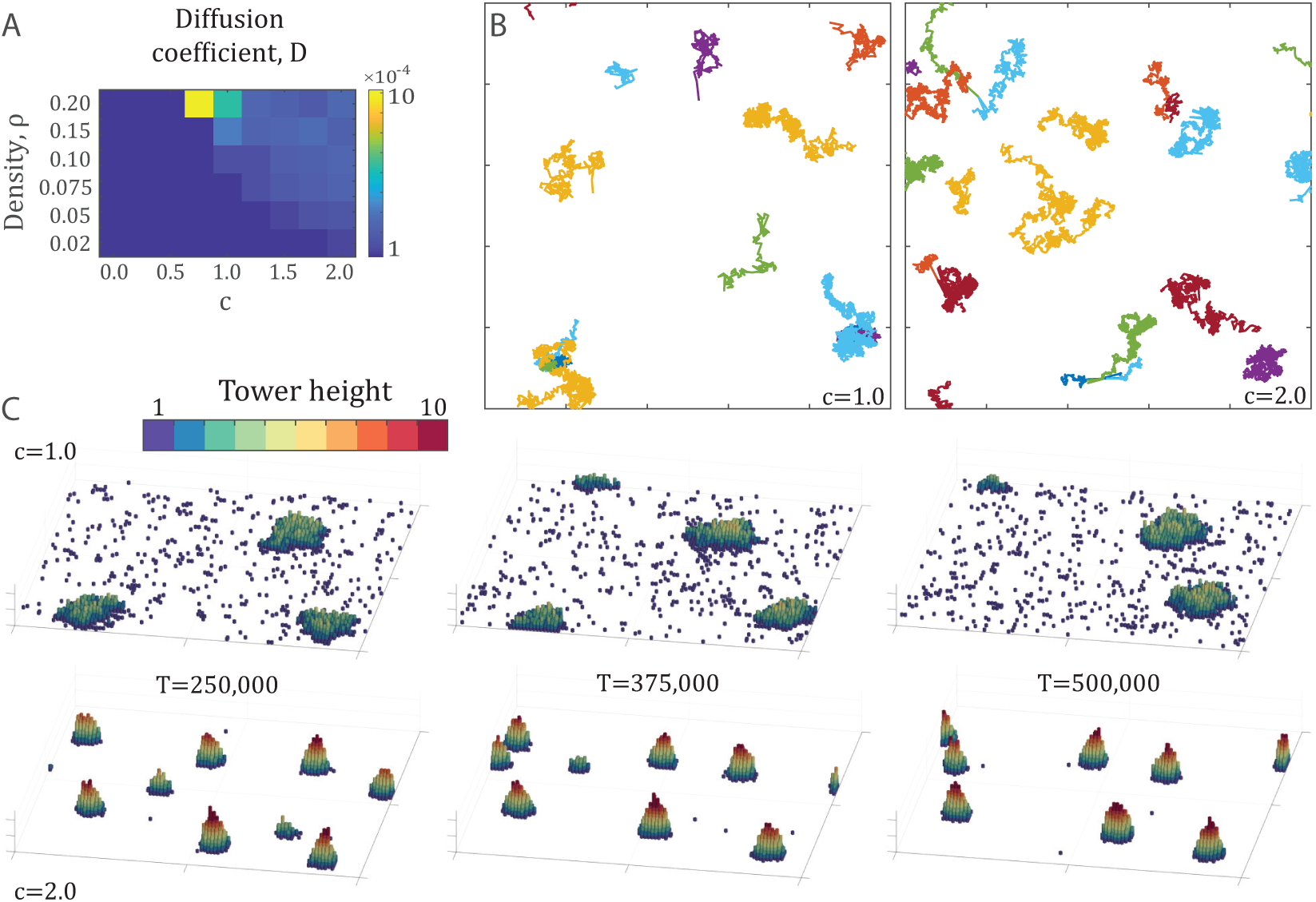
(A) Heat map of diffusion coefficients across the density-attraction space of Figure 4. Each point represents the mean diffusion coefficient (calculated by (3) of all tower trajectories over ten simulations of 500,000 time steps. The diffusion coefficient is calculated over the first 37,500 time steps of each trajectory. (B) Example trajectories from two simulations at the density *ρ* = 0.2 with attraction factors *c* = {1.0, 2.0}, with points every 250 time steps for 500,000 time steps. (C) Snapshots in time of the same simulations as in the top-right, showing the motion, shape change, and appearance and disappearance of towers over time. Each panel shows the entire 100 × 100 arena.

**Figure 6:**
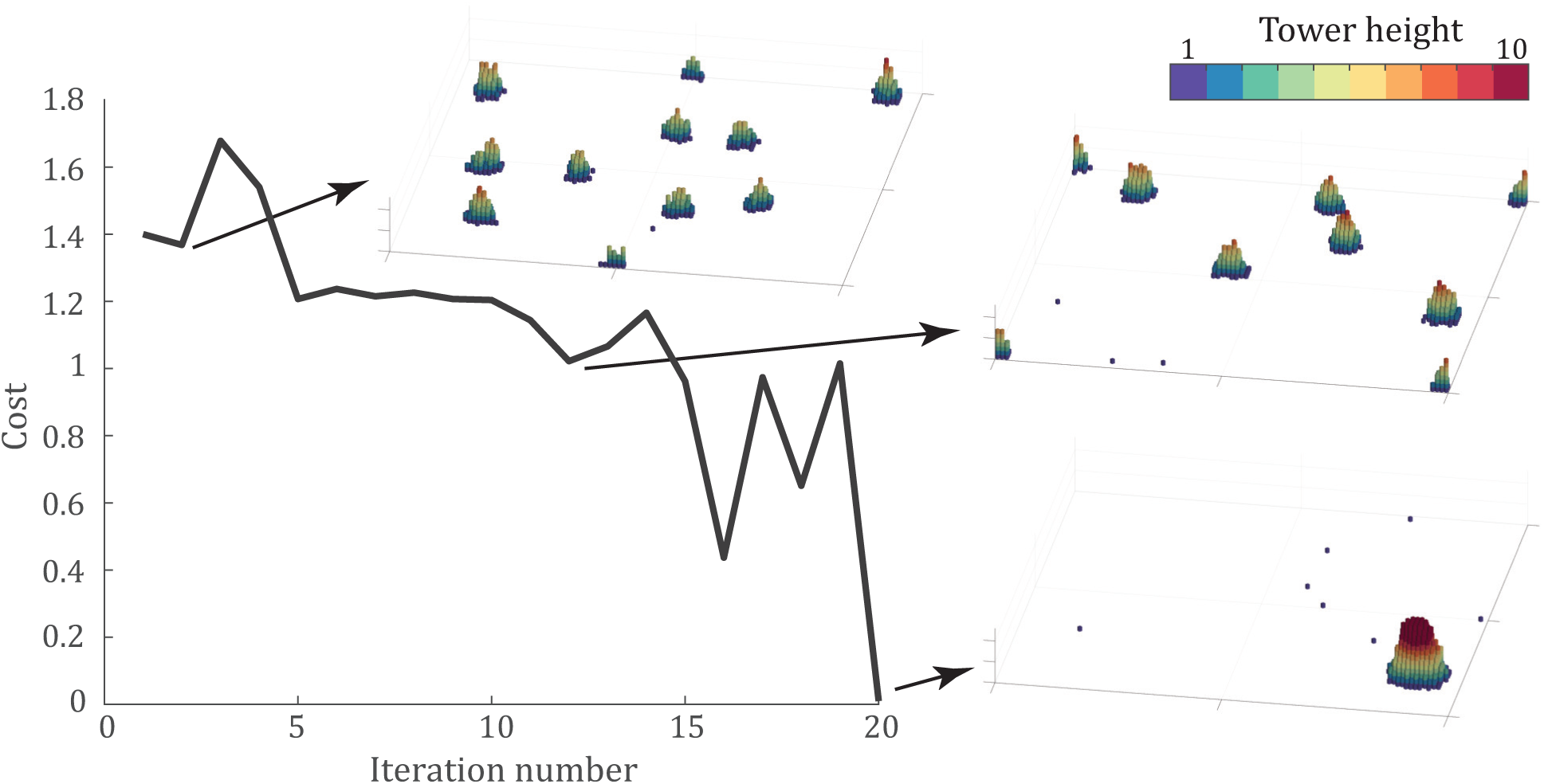
Optimization of the cost function given by (4) using the CMA-ES algorithm [Hansen et al., 2003]. Panels show the final configuration of the best simulation at iterations 2, 12, and 20 after 50,000 time steps. Each panel shows the entire 100 × 100 arena. The algorithm converges in 20 iterations, to the parameter set *P*_*u*_ = 0.938, *k*_*nl*_ = 0.029, and *c* = 2.56, with the optimal configuration shown in the bottom-right.

### 3.5 Tower optimization

One question that still remains is, what parameter values are optimal for tower building? To answer this, we need to think about what may constitute optimal. It may be that the optimal tower reaches as high as possible, which would, in practice, allow as many agents as possible to attach to a support structure. Or, for robotics applications, this would allow the tower to reach higher heights. On the other hand, it may be best to include as many individuals as possible in the tower, and the optimal tower would be the one that includes every single agent in the tower. As observed in Phonekeo et al. [2017], fire ants built towers that equally distribute load among the individuals. Therefore, an optimal tower from their perspective may be one that optimizes for load distribution. In this section, we use a genetic algorithm to explore optimal tower building considering each of these optimization targets.

To search for an optimal tower, we employ the Covariance Matrix Adaptation - Evolutionary Strategy (CMA-ES) algorithm developed by Hansen et al. [2003]. This algorithm randomly generates parameter sets within the search space and evaluates a cost function for each parameter set. From the results of this function evaluation, it updates the covariance matrix to expand in the direction of the most optimal value. Using the updated covariance matrix, the algorithm generates new parameter sets and repeats the process until convergence, generally defined as finding a parameter set with a cost function below some threshold.

We applied the CMA-ES algorithm to the tower-building model introduced above, using the average final properties of 3 trials for each parameter set across 10 parameter sets per iteration. For the optimization, we choose a cost function defining the optimal tower as the largest tower, both in terms of tower height and number of individuals within the tower. Therefore, the cost function is given by,

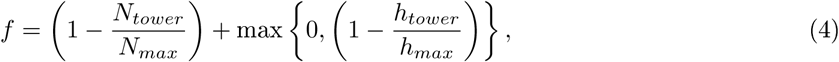

where *N*_*tower*_ and *h*_*tower*_ represent the number of individuals and height of the largest tower, *N*_*max*_ is the number of individuals in the simulation, and *h*_*max*_ is a prescribed maximum height. The height term is included to ensure that the results are effectively tower-like, preventing the optimal tower from simply achieving a large, wide aggregation. From the results of the attraction sweep in Figure 3A, we observe that *h*_*max*_ = 14 is an approximate upper bound on tower height, so it is therefore chosen as *h*_*max*_ for the purpose of this optimization. Note that a tower height of *h*_*tower*_ ≥ *h*_*max*_ results in a zero second term, and the simulation therefore allows for a taller tower.For the purposes of optimization, we reduce the simulation time to 50,000 time steps. This serves the practical role of making iterated simulation possible, but also places an effective minimization of convergence time. Therefore, we are optimizing for a tower that maximizes both height and number of agents quickly (within 50,000 time steps).

Figure 6 shows the progression of the minimum cost at each iteration of the CMA-ES algorithm along with snapshots of intermediate results to show the algorithm’s progress. The optimal tower occurs for the parameters, *P*_*u*_ = 0.938, *k*_*nl*_ = 0.029, and *c* = 2.56, which led to a tower of 993 agents reaching 16 agents tall after 50,000 time steps. The final cost function, averaged over three trials, was *f* = 0.01. A video of one simulation with these parameters is shown in supplementary video S7.

The CMA-ES optimization code of Hansen et al. [2003] applied to the present model may be found in the supplementary material (S8) to allow future research and consideration of other conditions of optimal tower-building. For example, when designing a robotic system where each individual robot has a maximum load capability, it may be necessary to calculate the maximum load experienced by an individual in the tower and add that term to cost function.

## 4 Discussion

In this work, we have extended a previously proposed set of local rules to replicate the tower-building behavior of red imported fire ants, *Solenopsis invicta*. This model and its insights will allow for the design of control strategies for tower-building swarm robotics and greater insight into the collective behavior of social insects. The results presented above show that individuals moving under the influence of local attraction are able to form large towers. We find that an attractive force is necessary for significant tower-building and show the impacts of this attractive force over a range of locking and unlocking parameters as well as a range of densities. We find that the system contains a sudden phase transition as the attraction parameter is varied, and that this phase transition is density-dependent. Finally, the largest towers, in both height and number of individuals, occur with a combination of very strong attraction and highly probable unlocking.

On the other hand, without attraction, no towers form, as shown in the *c* = 0 case of Figure 2 and supplemental video S2 and discussed further in the Appendix A and Figure A1. The effective force of attraction may also be thought of as a desire of the ants to climb, because the tallest available square to move toward will also have the most neighbors.

Near the phase transition, a critical slowing down occurs, and there are parameter sets that do not result in tower formation within a simulation time of 500,000 time steps. This critical slowing down is reminiscent of other examples of systems with phase transitions, such as the spin-glass model, the Ising model, and molecular dynamics models [Dasgupta et al., 1979, Hu, 2013]. Further from the phase transition (*c* ≫ *c*^∗^), towers form rapidly, but the possibility exists for these towers to encounter one another and merge into larger towers. The number of individuals in the tower and tower motion are largest just after the phase transition, but the largest height occurs with stronger attraction.

Our results also illustrate the exploration-exploitation trade-off, which balances attraction forces with random movement and unlocking events. Following this trade-off, stronger attraction may lead to higher towers with fewer individuals, as the attraction rapidly draws individuals from the edge of the aggregation toward the center of the tower, and therefore upward. This balance of unlock probability and attraction is found through the combined optimization of number of individuals and tower height, which discovered that with an unlock probability of *P*_*u*_ = 0.938 and an attraction of *c* = 2.56, it is possible to include nearly all of the individuals in a simulation, with a tower reaching a height of 16 layers. This large unlock probability of the largest towers in our simulations connects with the observation from Phonekeo et al. [2017] that, in the experimental system, the fire ants are constantly rebuilding their tower and circulating ants throughout the tower.The work of Phonekeo et al. [2017] showed that fire ants build towers of constant load, and future optimization work could incorporate the load experienced by each individual to achieve towers that prioritize stability.

The results of the parameter sweep in density values showed both similarities and differences across densities. In general, for a fixed attraction ratio *c*, the tower height-diameter ratio remains fairly constant, even as the numbers of agents per tower and tower height vary. The biggest difference across densities is the change in critical attraction parameter, *c*^∗^. These observations lead to testable hypotheses for animal experiments. Below a certain density threshold, tower formation should cease, due to the move past the critical attraction. Additionally, the height-diameter ratio should remain constant across a large range of densities. Finally, we have shown that the towers built in our simulations move over time, with a diffusion coefficient that is dependent on both attraction and density, and should be taken into account when considering practical application to robotics.

This work also lays the groundwork for future robotic studies, where robots are able to built towers out of themselves in a manner similar to, for example, the M-blocks of Romanishin et al. [2015]. The tower is a ubiquitous structure in building, and designing rigorous control strategies for tower-building represents a fundamental starting point toward fully autonomous, locally-sensed swarm building applications. In practice, a tower of robots could be useful in the case of, for example, seeing over obstacles, providing scaffolding for climbing, or clearly marking a location of interest. The control strategies introduced in the present study could be further modified to more closely replicate experimentally-observed fire ant behavior, developing a control strategy for interacting with a support structure.

## Author Contributions

NM, JC, TS, and JL developed the initial version of the present model as a project in a course taught by OP. GN refined the model, conducted simulations, and wrote and edited the manuscript. OP supervised the research and edited the manuscript.

## Acknowledgments

We thank Prof. David L. Hu, and members of the Peleg lab for insightful discussions, and the BioFrontiers Institute for the utmost support.

## Supplementary Material

**Supplementary Video S1.** The diffusion-limited aggregation case of the model, with *P*_*u*_ = 0, *k*_*nl*_ = 1, *c* = 0. The video is shown at a speed of 120 time steps per second.

**Supplementary Video S2.** Simulation in which no aggregations form, with The *P*_*u*_ = 0.2, 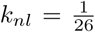, *c* = 0. The video is shown at a speed of 10,000 time steps per second.

**Supplementary Video S3.** Simulation in which large, wide aggregations form, with *P*_*u*_ = 0.2, 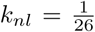, *c* = 1. The video is shown at a speed of 10,000 time steps per second.

**Supplementary Video S4.** Simulation in which many steep towers form, with *P*_*u*_ = 0.2, 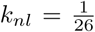, *c* = 2. The video is shown at a speed of 10,000 time steps per second.

**Supplementary Video S5.** Simulation with large, wide moving aggregations in a dense environment, with *P*_*u*_ = 0.2, 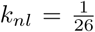, *c* = 1. and *N* = 2,000 individuals. The video is shown at a speed of 10,000 time steps per second.

**Supplementary Video S6.** Simulation with many steep moving aggregations in a dense environment, with *P*_*u*_ = 0.2, 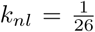, *c* = 2. and *N* = 2,000 individuals. The video is shown at a speed of 10,000 time steps per second.

**Supplementary Video S7.** Simulation of the results of tower optimization, with *P*_*u*_ = 0.938, *k*_*nl*_ = 0.029, *c* = 2.56. The video is shown at a speed of 2,500 time steps per second.

**Supplementary Code S8.** Three MATLAB code files, included in a. zip file. TowerSimulation.m provides a function to run a single simulation, TowerAnalysis.m provides the analysis of the resulting towers, and TowerOptimization.m provides the CMA-ES code used in the present study to optimize model parameters. Supplemental code not uploaded as part of BioRxiv submission, if you would like access to Supplemental Code.

## A Model results in the absence of attraction, *c* = 0

In the absence of attraction, significant tower formation does not occur, even with relaxed locking and unlocking probabilities. This is discussed above in Section 2.2.2, but we will discuss it further here. Figure A1 shows the effects of unlocking or neighbor-influenced lock probability, with the pure DLA case shown in the boxed-in frame.

The horizontal axis of Figure A1 shows the effects of unlocking: small values of unlocking allow initially fractal structures to break up into smaller aggregations, but too much unlocking leads to no aggregations at all. The effect of decreasing locking factor *k*_*nl*_ alone is shown in the left column. This rule modification alone does allow rounder aggregations to form, but the aggregations remain shallow. Without unlocking, it is unlikely that new agents make their way toward the center of the aggregation.

The effects of combining the two rule modifications are discussed above and shown most clearly in the *c* = 0 panels of Fig. 2. No combination of locking and unlocking parameters leads to consistent tower formation. The maximum aggregation size is only 150 individuals and no tower height exceeds 2 agents high. Without attraction, small aggregations are able to form, but none of them come close to approximating the tower formation observed in Figure 1A or those described in Phonekeo et al. [2017].

## B Supplementary model definitions

In this section, we provide equations described in the text.

In Section 2, we describe the periodic boundary condition. An agent’s position at time step *t* + 1, under the periodic boundary condition, is given by,

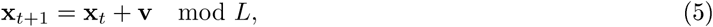

where **x**_*t*_ and **v**_*t*_ are its position and velocity at time step *t*. Periodic boundaries are also taken into account when calculating distances between agents, with the distance between positions **x**_1_ and **x**_2_ defined by,

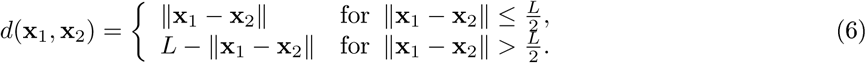

Any two points with a distance further than 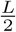 are closer to the border, and their distance is therefore *L* − ‖**x**_1_ − **x**_2_‖.

The normalized velocity 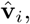 which defines the agent’s motion in Section 2.2.3, is given by,

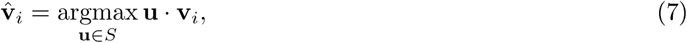

where *S* is the set of velocities that reach adjacent pixels in one time step,

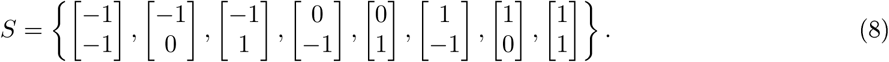

**Figure A1:**
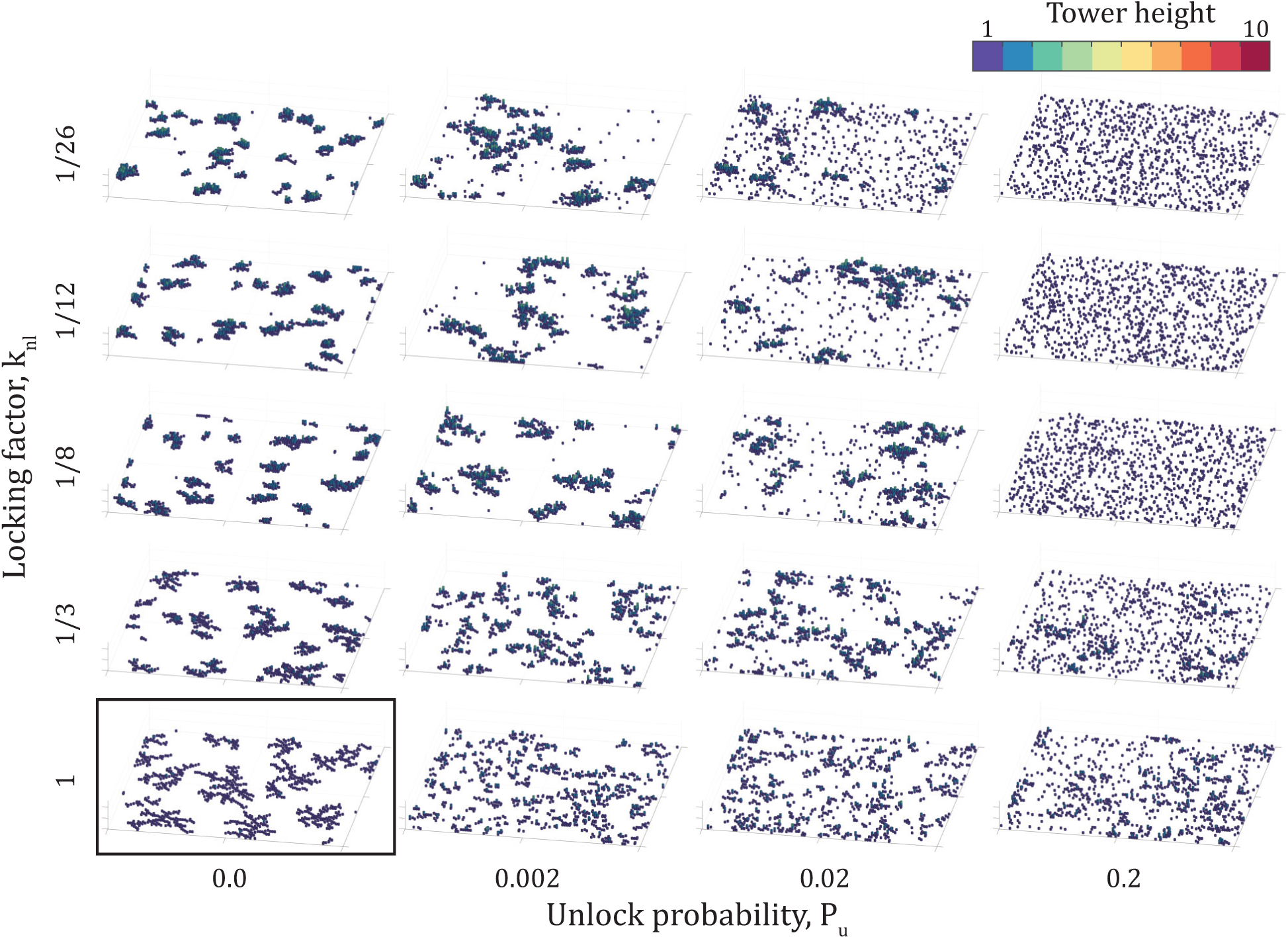
Figure A1: Final configuration of simulations, after 500,000 time steps, incorporating only the effect of unlock probability *P*_*u*_ and neighbor-influenced lock factor *k*_*nl*_. Each panel shows the entire 100 × 100 arena. The panel surrounded by the box represents the case of no rule modifications, leading to diffusion-limited aggregation.

The normalized velocity 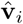 is the adjacent pixel which has the direction closest to the velocity **v**_*i*_ defined by (2).

## References

Jean-Marc Ame, Colette Rivault, and Jean-Louis Deneubourg. Cockroach aggregation based on strain odour recognition. Animal Behaviour, 68(4):793–801, 2004. ISSN 0003-3472. doi: https://doi.org/10.1016/j.anbehav.2004.01.009. URL http://www.sciencedirect.com/science/article/pii/S0003347204002210.

Carl Anderson, Guy Theraulaz, and J.-L. Deneubourg. Self-assemblages in insect societies. Insectes sociaux, 49(2):99–110, 2002.

P. Bak. How nature works: the science of self-organized criticality. Springer, 1996.

Eric Bonabeau, Sylvain Guérin, Dominique Snyers, Pascale Kuntz, and Guy Theraulaz. Three-dimensional architectures grown by simple ‘stigmergic’ agents. BioSystems, 56(1):13–32, 2000.

A Buffin and SC Pratt. Cooperative transport by the ant novomessor cockerelli. Insectes sociaux, 63(3): 429–438, 2016.

Erol Şahin, Thomas H Labella, Vito Trianni, J-L Deneubourg, Philip Rasse, Dario Floreano, Luca Gambardella, Francesco Mondada, Stefano Nolfi, and Marco Dorigo. Swarm-bot: Pattern formation in a swarm of self-assembling mobile robots. In IEEE International Conference on Systems Man and Cybernetics, volume 4, pages 6-pp. IEEE, 2002.

Chandan Dasgupta, Shang-keng Ma, and Chin-Kun Hu. Dynamic properties of a spin-glass model at low temperatures. Physical Review B, 20(9):3837, 1979.

Emanuela Del Dottore, Ali Sadeghi, Alessio Mondini, Virgilio Mattoli, and Barbara Mazzolai. Toward growing robots: a historical evolution from cellular to plant-inspired robotics. Frontiers in Robotics and AI, 5:16, 2018.

J. L. Deneubourg, J. C. Grégoire, and E. Le Fort. Kinetics of larval gregarious behavior in the bark beetledendroctonus micans (coleoptera: Scolytidae). Journal of Insect Behavior, 3(2):169–182, Mar 1990. ISSN 1572-8889. doi: 10.1007/BF01417910. URL https://doi.org/10.1007/BF01417910

J. L. Deneubourg, A. Lioni, and C. Detrain. Dynamics of aggregation and emergence of cooperation. The Biological Bulletin, 202(3):262–267, 2002. doi: 10.2307/1543477. URL https://doi.org/10.2307/1543477. PMID: 12086998.

Nigel R. Franks. Thermoregulation in army ant bivouacs. Physiological entomology, 14(4):397–404, 1989.

Roderich Groẞ, Elio Tuci, Marco Dorigo, Michael Bonani, and Francesco Mondada. Object transport by modular robots that self-assemble. In Proceedings 2006 IEEE International Conference on Robotics and Automation 2006. ICRA 2006., pages 2558–2564. IEEE, 2006.

Heiko Hamann. Swarm robotics: A formal approach. Berlin: Springer, 2018.

Nikolaus Hansen, Sibylle D Müller, and Petros Koumoutsakos. Reducing the time complexity of the derandomized evolution strategy with covariance matrix adaptation (cma-es). Evolutionary computation, 11(1): 1–18, 2003.

Chin-Kun Hu. Slow dynamics in proteins and polymer chains. In AIP Conference Proceedings, volume 1518, pages 541–550. AIP, 2013.

Dana Hughes, Christoffer Heckman, and Nikolaus Correll. Materials that make robots smart. The International Journal of Robotics Research, page 0278364919856099, 2019.

Raphael Jeanson, Colette Rivault, Jean-Louis Deneubourg, Stephane Blanco, Richard Fournier, Christian Jost, and Guy Theraulaz. Self-organized aggregation in cockroaches. Animal Behaviour, 69(1):169–180, 2005. ISSN 0003-3472. doi: https://doi.org/10.1016/j.anbehav.2004.02.009. URL http://www.sciencedirect.com/science/article/pii/S0003347204002428.

Gerald Kastberger, Frank Weihmann, and Thomas Hoetzl. Self-assembly processes in honeybees: the phenomenon of shimmering. In Honeybees of Asia, pages 397–443. Springer, 2011.

Gerald Kerth. Causes and consequences of sociality in bats. Bioscience, 58(8):737–746, 2008.

Herbert Levine, Lev Tsimring, and David Kessler. Computational modeling of mound development in dictyostelium. Physica D: Nonlinear Phenomena, 106(3-4):375–388, 1997.

Nathan J. Mlot, Craig A. Tovey, and David L. Hu. Fire ants self-assemble into waterproof rafts to survive floods. Proceedings of the National Academy of Sciences, 108(19):7669–7673, 2011.

Nathan J. Mlot, Craig Tovey, and David L. Hu. Dynamics and shape of large fire ant rafts. Communicative & integrative biology, 5(6):590–597, 2012.

Ricard Matas Navarro and Suzanne M Fielding. Clustering and phase behaviour of attractive active particles with hydrodynamics. Soft Matter, 11(38):7525–7546, 2015.

AD Panagiotou, MW Curtin, H Toki, DK Scott, and PJ Siemens. Experimental evidence for a liquid-gas phase transition in nuclear systems. Physical Review Letters, 52(7):496, 1984.

Sulisay Phonekeo, Nathan Mlot, Daria Monaenkova, David L. Hu, and Craig Tovey. Fire ants perpetually rebuild sinking towers. Royal Society open science, 4(7):170475, 2017.

Sven Reynaert, Paula Moldenaers, and Jan Vermant. Control over colloidal aggregation in monolayers of latex particles at the oil-water interface. Langmuir, 22(11):4936–4945, 2006a. doi: 10.1021/la060052n. URL https://doi.org/10.1021/la060052n. PMID: 16700578.

Sven Reynaert, Paula Moldenaers, and Jan Vermant. Control over colloidal aggregation in monolayers of latex particles at the oil-water interface. Langmuir, 22(11):4936–4945, 2006b.

Craig W. Reynolds. Flocks, herds and schools: A distributed behavioral model. In ACM SIGGRAPH computer graphics, volume 21, pages 25–34. ACM, 1987.

John W Romanishin, Kyle Gilpin, Sebastian Claici, and Daniela Rus. 3d m-blocks: Self-reconfiguring robots capable of locomotion via pivoting in three dimensions. In 2015 IEEE International Conference on Robotics and Automation (ICRA), pages 1925–1932. IEEE, 2015.

M Rovere, DW Heermann, and K Binder. The gas-liquid transition of the two-dimensional lennard-jones fluid. Journal of Physics: Condensed Matter, 2(33):7009, 1990.

Roald C Roverud and Mark A Chappell. Energetic and thermoregulatory aspects of clustering behavior in the neotropical bat noctilio albiventris. Physiological Zoology, 64(6):1527–1541, 1991.

Erol Şahin. Swarm robotics: From sources of inspiration to domains of application. In International workshop on swarm robotics, pages 10–20. Springer, 2004.

Thomas D Seeley. Honeybee democracy. Princeton University Press, 2010.

Linda G Shapiro. Connected component labeling and adjacency graph construction. In T. Yung Kong and Azriel Rosenfeld, editors, Topological Algorithms for Digital Image Processing, volume 19 of Machine Intelligence and Pattern Recognition, pages 1–30. North-Holland, 1996.

Petras Swissler and Michael Rubenstein. Fireant: A modular robot with full-body continuous docks. In 2018 IEEE International Conference on Robotics and Automation (ICRA), pages 6812–6817. IEEE, 2018.

Guy Theraulaz and Eric Bonabeau. Modelling the collective building of complex architectures in social insects with lattice swarms. Journal of Theoretical Biology, 177(4):381–400, 1995.

Guy Theraulaz, Eric Bonabeau, Stamatios C. Nicolis, Ricard V. Sole, Vincent Fourcassie, Stephane Blanco, Richard Fournier, Jean-Louis Joly, Pau Fernández, Anne Grimal, Patrice Dalle, and Jean-Louis Deneubourg. Spatial patterns in ant colonies. Proceedings of the National Academy of Sciences, 99(15): 9645–9649, 2002. ISSN 0027-8424. doi: 10.1073/pnas.152302199. URL https://www.pnas.org/content/99/15/9645.

T Umeda and K Inouye. Theoretical model for morphogenesis and cell sorting in dictyostelium discoideum. Physica D: Nonlinear Phenomena, 126(3-4):189–200, 1999.

FJ Vernerey, E Benet, L Blue, AK Fajrial, S Lalitha Sridhar, J Lum, G Shakya, KH Song, AN Thomas, and MA Borden. Biological active matter aggregates: Inspiration for smart colloidal materials. Advances in colloid and interface science, 2018.

Tamas Vicsek, András Czirók, Eshel Ben-Jacob, Inon Cohen, and Ofer Shochet. Novel type of phase transition in a system of self-driven particles. Physical review letters, 75(6):1226, 1995.

Valeriy A. Vlasov. Activated complex theory of nucleation. The European Physical Journal E, 42(3):36, Mar 2019. ISSN 1292-895X. doi: 10.1140/epje/i2019-11797-7. URL https://doi.org/10.1140/epje/i2019-11797-7

P. W. Voorhees. The theory of ostwald ripening. Journal of Statistical Physics, 38(1):231–252, Jan 1985. ISSN 1572-9613. doi: 10.1007/BF01017860. URL https://doi.org/10.1007/BF01017860.

Aaron Waters, Fran;ois Blanchette, and Arnold D Kim. Modeling huddling penguins. PLoS One, 7(11): e50277, 2012.

Justin Werfel, Kirstin Petersen, and Radhika Nagpal. Designing collective behavior in a termite-inspired robot construction team. Science, 343(6172):754–758, 2014.

T.A. Witten Jr., and Leonard M. Sander. Diffusion-limited aggregation, a kinetic critical phenomenon. Physical review letters, 47(19):1400, 1981.

